# Estimates of persistent inward currents in lower limb muscles are not different between inactive, resistance-trained and endurance-trained young individuals

**DOI:** 10.1101/2023.07.13.548353

**Authors:** Valentin Goreau, François Hug, Anthony Jannou, François Dernoncourt, Marion Crouzier, Thomas Cattagni

**Affiliations:** Movement-Interactions-Performance, MIP, UR 4334, Nantes Université, Nantes, France; LAMHESS, Université Côte d’Azur, Nice, France; Department of Movement Sciences, Leuven, Belgium

**Keywords:** Neuromodulation, physical training, motor units, motoneuron, electromyography

## Abstract

Persistent inward currents (PICs) increase the intrinsic excitability of α-motoneurons. The main objective of this study was to determine whether estimates of α-motoneuronal PIC magnitude is influenced by chronic endurance and resistance training. We also aimed to investigate whether there is a relationship in the estimates of α-motoneuronal PIC magnitude between muscles. Estimates of PIC magnitude were obtained in three groups of young individuals: resistance-trained (n=12), endurance-trained (n=12), and inactive (n=13). We recorded high-density surface electromyography (HDsEMG) signals from tibialis anterior, gastrocnemius medialis, soleus, vastus medialis, and vastus lateralis. Then, signals were decomposed with convolutive blind source separation to identify motor units spike trains. Participants performed triangular isometric contractions to a peak of 20% of their maximum voluntary contraction. A paired-motor-unit analysis was used to calculate ΔF, which is assumed to be proportional to PIC magnitude. Despite the substantial differences in physical training experience between groups, we found no differences in ΔF, regardless of the muscle. Significant correlations of estimates of PICs magnitude were found between muscles of the same group (VL-VM, SOL-GM). Only one correlation (out of 8) between muscles of different groups was found (GM and TA). Overall, our findings suggest that estimates of PIC magnitude in the lower limb muscles are not influenced by physical training experience in healthy young individuals. They also suggest muscle-specific and muscle group-specific regulations of the diffuse monoamine inputs.

## INTRODUCTION

There is extensive evidence that endurance and resistance training can elicit specific adaptations in motor unit (MU) activity during submaximal contractions. Specifically, when considering young healthy adults, endurance training has been associated with a lower MU discharge rate (Vila-Chã et al., 2010; Dimmick et al., 2018; Trevino et al., 2022), and resistance training has been associated with an increase in MU discharge rate (Del Vecchio et al., 2019). Previous work has primarily focused on examining the ionotropic inputs to MUs to explain these training-induced changes (Škarabot et al., 2021). For example, studies using transcranial magnetic stimulation observed corticospinal adaptations after resistance training (Siddique et al., 2020), whereas no such adaptations have been observed after endurance training (Millet & Temesi, 2019). The role of neuromodulatory inputs in the adaptation of MU discharge characteristics with chronic endurance and resistance training remains to be explored.

Neuromodulatory inputs are promoted by monoamines such as serotonin (5-HT), that bind to G-proteins and activate voltage-dependent channels on the motoneuron dendrites (Heckman, Johnson *et al*., 2008). Notably, neuromodulatory inputs lead to the generation of strong persistent inward currents (PICs), which are depolarizing currents generated by voltage-sensitive sodium and calcium channels. These currents have the ability to increase the intrinsic excitability of the target motoneuron (Gorassini *et al*., 2002; Heckman *et al*., 2005). This increase in intrinsic excitability results in a substantial amplification of the input-output gain of α-motoneurons up to fivefold (Lee & Heckman, 2000*a*).

Together with an increase in MU discharge rate and force production, an increase in estimates of PIC magnitude have been reported after 6 weeks of resistance training in the soleus muscle of older adults (Orssatto *et al*., 2023). It echoes previous work conducted in a rat model, which reported an increase in intrinsic excitability characterized by lower input currents required to achieve rhythmic discharge (Krutki *et al*., 2017). Of note, a similar increase in excitability has been observed after endurance training in rats (MacDonell & Gardiner, 2018). In addition, animal studies have demonstrated that chronic treadmill exercises enhance the activation of serotonin neurons in the raphe nuclei (Ji *et al*., 2017; Ge & Dai, 2020). This could lead to an increase in serotonin release, which, in turn, may promote α-motoneuronal PICs. Together, these results suggest that both resistance and endurance training could promote α-motoneuronal PICs in healthy young individuals.

Descending monoaminergic inputs to α-motoneurons are diffuse, simultaneously influencing multiple motor pools within an individual (Heckman, Hyngstrom *et al*., 2008; Wei *et al*., 2014). This likely explains why a 30-second handgrip contraction at 40% of maximal voluntary contraction (MVC) increased estimates of PIC magnitude in distant muscles, i.e., soleus and tibialis anterior (Orssatto *et al*., 2022). These results suggest a relationship in PIC magnitude across muscles, the overall level of descending monoamines leading to individual PIC magnitude profiles (e.g., individuals having consistently high or low PIC magnitude across all muscles). In other words, PIC magnitude of α-motoneurons would be positively correlated across muscles, but this remains to be confirmed.

The main objective of this study was to determine whether estimates of α-motoneuronal PIC magnitude are influenced by endurance and resistance training. We also aimed to investigate whether there is a relationship between the estimates of α-motoneuronal PIC magnitude of different lower limb muscles. To this end, we compared three groups of participants with different physical training experience: i) resistance training, ii) endurance training, and iii) inactive. MU activity was extracted using high-density surface EMG (HDsEMG). Triangular contractions were used to estimate α-motoneuronal PICs using the paired-MU technique (Gorassini et al. 2002; Hassan et al. 2020), and to investigate peak discharge rates. We hypothesised that estimates of PIC magnitude would be higher in both the resistance and endurance training groups compared to the inactive group, irrespective of the muscle. No hypothesis was established with respect to differences between training groups. Additionally, we hypothesised that the estimates of PIC magnitude would be positively correlated across muscles, suggesting the existence of individual PIC magnitude profiles.

## METHODS

### Participants

Thirty-seven males aged between 18 and 35 years old, were recruited according to their physical training experience (at least one year). They were divided into either a resistance-trained group (n=12), an endurance-trained group (n=12) and an inactive group (n=13). Physical training experience was assessed using the ONAPS-PAQ questionnaire, which provides a standardised assessment of the type and volume of physical activity in the previous year. Detailed characteristics of the participants are presented in Table 1.

**Table 1:**
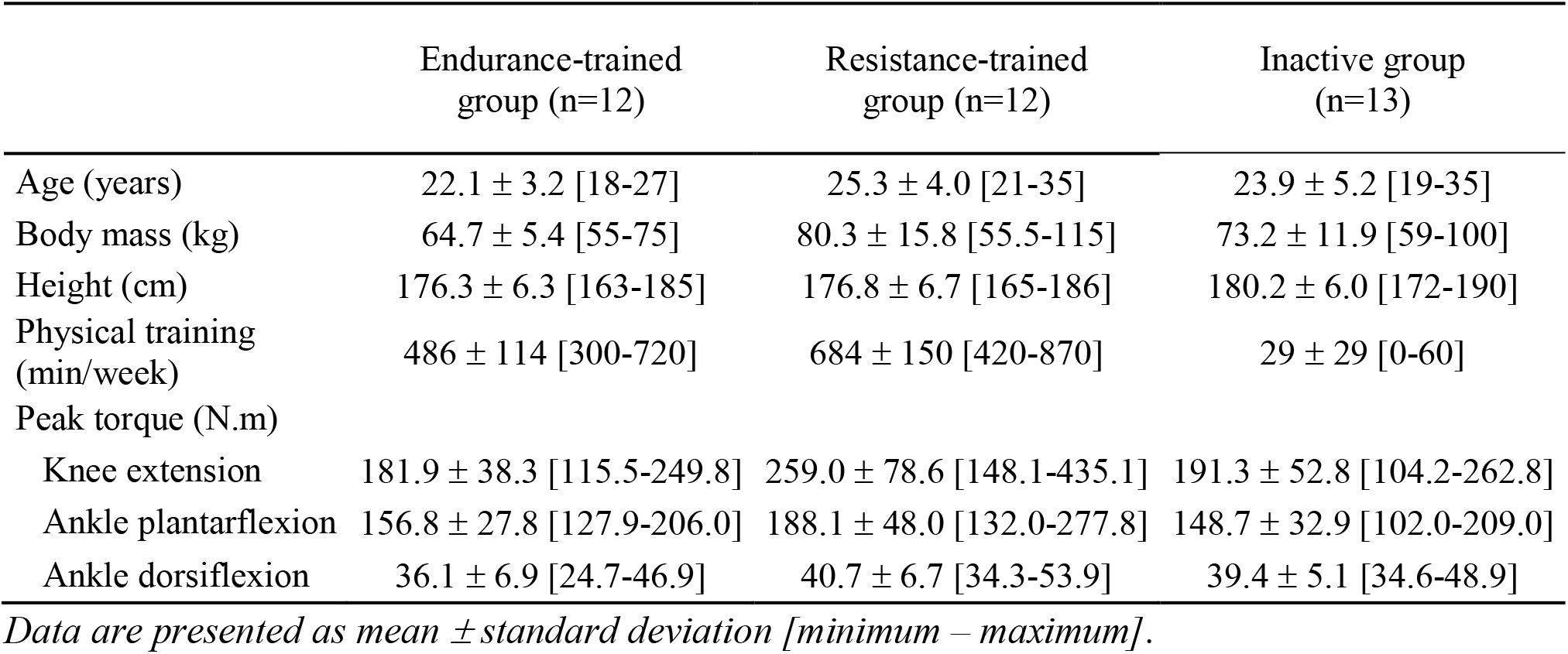
Participant characteristics.

Participants of the resistance group performed resistance-based training on average 684 ± 150 min/week, which included training of the lower limb muscles. Main disciplines included powerlifting (n=2), bodybuilding (n=1) and weightlifting (n=9). Participants of the endurance group performed endurance-based training on average 486 ± 114 min/week consisting of triathlon (n=4), running (n=6), cycling (n=1) and duathlon (n=1). Participants in the trained groups could engage in other training, for example, endurance training for resistance-trained individuals and vice versa, for no more than 30 min per week. Participants in the inactive group participated in a physical training activity on average 29 ± 29 min/week. All individuals in the inactive group performed less than the recommended cut-off of 150 min/week of moderate physical activity or 75 min/week of vigorous physical activity or a combination of both (Bull et al., 2020).

Participants were required to report no lower limb injuries in the preceding 6 months. Participants were instructed not to consume any caffeine (e.g. coffee) or engage in lower limb training for 24 hours prior to the evaluation. The study protocol was approved by the local ethics committee (CERNI, Nantes Université, IRB00013074). The experimental protocol and any potential discomfort were explained to participants and written informed consent was obtained prior to testing.

### Experimental protocol

Evaluations were performed in five muscles of the participant’s dominant leg (used to kick a ball): soleus (SOL) and gastrocnemius medialis (GM) during plantar flexion, tibialis anterior (TA) during dorsiflexion, vastus medialis (VM) and vastus lateralis (VL) during knee extension. These three different tasks were performed in a randomised order, with a similar experimental procedure. Participants first performed a standardized warm-up, which consisted of three 5-second contractions at each of the following intensities: 30%, 60%, and 90% of their subjectively estimated maximal MVC. There was a 20-s rest period between each contraction.

#### Positions for testing the different muscle groups

For plantar flexion and dorsiflexion, participants were tested in a seated position with the hip angle at 80° (neutral position=0°), the knee angle at 0° (neutral position), and the ankle angle at 10° of plantar flexion (neutral ankle position). The lateral malleolus was aligned with the axis of rotation of the dynamometer (Biodex System 4, Biodex Medical system, Shirley, NY). The foot, trunk, and thigh were securely strapped to the dynamometer. Participants were asked to cross their arms across their chest during each contraction.

For knee extension, participants were tested in a seated position with the hip angle at 80° and the knee joint angle at 90°. The lateral epicondyle of the femur was aligned with the axis of rotation of the dynamometer. The strap was positioned just above the malleoli. The trunk and thigh were securely strapped to the dynamometer and participants were asked to cross their arms across their chest during each contraction.

#### Isometric maximal voluntary contraction

After a 2-min rest period, participants performed three 4-5 s MVCs separated by a 2-min rest period. They received real-time visual feedback of the torque signal during each effort and strong verbal encouragements were provided.

#### Isometric triangular voluntary contractions

Triangular contractions to 20% of MVC torque were used to calculate ΔF using the paired-MU technique (Gorassini *et al*., 2002). These contractions lasted 20 s and consisted of a 10-s ramp-up phase and a 10-s ramp-down phase. Participants were instructed to move a real-time torque feedback displayed on a computer screen as close as possible to a trace representing the 20% triangular contraction. Participants were first familiarised with the task. Once they were able to accurately follow the torque feedback, a 5-min rest period was respected. Data were collected on two trials of triangular contractions with a 3-min rest interval between each. Trials were excluded and repeated if the torque trajectory was not closely matched, for example a sharp increase or decrease in torque signal.

### Data recordings

#### Electromyography recordings

HDsEMG signals were recorded from SOL and GM during plantar flexion, from TA during dorsiflexion, and from VM and VL during knee extension. A two-dimensional adhesive grid of 64 electrodes [13 × 5 gold-coated electrodes with one electrode absent on a corner; inter-electrode distance: 8 mm (ELSCH064NM2, OT Bioelettronica, Italy)] was placed over each muscle. The grids were aligned in the direction of the fascicles. Before mounting the electrode grids, the skin was shaved and then cleaned with an abrasive pad and alcohol. The adhesive grids were held on the skin using semi disposable biadhesive foam layers (SpesMedica, Battipaglia, Italy). The skin-electrode contact was made by filling the cavities of the adhesive layers with conductive paste (SpesMedica, Battipaglia, Italy). Reference electrodes (Kendall Medi-Trace, Canada) were placed over the patella. The EMG signals were recorded in monopolar mode, bandpass filtered (10–500 Hz), and digitised at a sampling rate of 2048 Hz using a multichannel HDsEMG acquisition system (EMG-Quattrocento, 400 channel EMG amplifier; OT Bioelettronica).

#### Mechanical recordings

Torque values for plantar flexors, dorsiflexors, and knee extensors were collected on the isokinetic dynamometer (Biodex System 3, Biodex Medical Systems, Shirley, NY, USA). The output signal from the dynamometer was collected at 2048 Hz using the multichannel HDsEMG acquisition system.

### Data analysis

#### HDsEMG decomposition

The HDsEMG signals were decomposed with the convolutive blind source separation method (Negro *et al*., 2016) implemented in a custom-made MATLAB (Version R2019b; The MathWorks Inc., Natick, MA) routine to iteratively identify and extract the sources of the interference HDsEMG signals, i.e., the MUs spike trains. After the automatic decomposition, all the MU spike trains were visually inspected and manually edited with the DEMUSE software (v5.01; The University of Maribor, Slovenia), as described previously (Del Vecchio *et al*., 2020; Hug *et al*., 2021). Only MUs with a pulse-to-noise ratio equal to or greater than 30 dB were kept for further analysis (Holobar *et al*., 2014). This decomposition procedure has been extensively validated using experimental and simulated signals (Holobar *et al*., 2014; Holobar & Farina, 2014). For each muscle, only the triangular contraction yielding the highest number of identified MUs was analysed. If both contractions presented the same number of identified MUs, the triangular contraction yielding the highest number of pairs of MUs was analyzed.

#### Estimating PIC magnitude and peak discharge rate

Instantaneous MU firing rates were calculated as the inverse of the interspike intervals of each MU spike train and smoothed using a 1-s Hanning window using a custom-written MATLAB script. The maximum value obtained from the Hanning filter of each MU was considered as the peak discharge rate. PIC magnitude was estimated using the paired MU analysis (Gorassini *et al*., 2002). MUs with a lower recruitment threshold (control units) were paired with MUs with a higher recruitment threshold (test units). ΔF was calculated as the change in discharge rates of the control MU from the moment of recruitment to the moment of de-recruitment of the test unit (Gorassini *et al*., 2002). In agreement with previous work (Hassan *et al*., 2020), criteria for inclusion of ΔF value from MUs pairs were as follows: (i) a test MU should discharge for at least 2 s, (ii) a test MU should be recruited at least 1 s after the control MU to ensure full activation of PIC, (iii) a test MU should be de-recruited at least 1.5 s prior to the control MU to prevent overestimation of ΔF, (iv) a coefficient of determination (*r*²) ≥ 0.7 should be observed between the smoothed discharge rate of the test and control MU (Kim *et al*., 2020). A high *r*² was chosen to ensure that the control and test units receive similar synaptic inputs, as several studies have reported the *r*² as a measure of common synaptic input between motor units (Gorassini *et al*., 2002; Powers *et al*., 2008; Stephenson & Maluf, 2011). As in previous studies, ΔF values are presented as ‘unit-wise’ averages for all suitable test–control unit pairs, thus reducing the number of ΔF values to one per test unit (Hassan *et al*., 2021; Khurram *et al*., 2021; Orssatto, Borg *et al*., 2021). When individual test units were paired with multiple control units that met inclusion criteria, the average ΔF value was obtained.

### Statistical analysis

The sample size for this study was based on the estimates of PICs (ΔF). The calculation was based on a one-way ANOVA analysis, assuming a standard deviation of 0.8 pulse per second (pps) in each group. A total sample size of 36 (12 in each group) was deemed necessary to achieve 80% power to detect a difference in ΔF of 0.8 pps between the groups with 5% significance level. This difference was based on a previous report of approximately 0.8 pps increase following 6 weeks of resistance training (Orssatto et al., 2023). One participant was added to the inactive group to compensate for missing data. Therefore, we recruited 37 participants.

All analyses were performed using R (v.4.1.1; R Core Team 2021, R Foundation for Statistical computing, Vienna, Austria). For descriptive analysis, mean and standard deviation values for each variable reported were calculated after averaging the data of the MUs for each muscle. The effects of physical activity on age, weight, height, time of physical training, knee extension MVC, plantar flexion MVC and dorsal flexion MVC were determined using separate one-way ANOVAs.

Further analyses were performed by considering all individual MUs instead of averaging them on a per-muscle basis. The effect of physical training experience and muscle on ΔF and peak discharge rate was assessed with separate linear mixed-effects models including physical training experience (endurance, resistance, inactive), muscle (TA, GM, SOL, VM, VL) and their interaction as fixed-effects. As random-effects, we included a random intercept for each individual as well as a random slope accounting for the muscle within each individual [e.g. ΔF ∼ muscle*group + (muscle | participant)]. Models were fitted using the lmerTest package (Kuznetsova *et al*., 2017). For all models, we confirmed that the assumptions of homoscedasticity, normality and independence of residuals were valid by graphical evaluations. The effect sizes derived from the F ratios were calculated with the partial omega-squared (ω2). When a significant effect was observed, Tukey post hoc correction was adopted for pairwise comparison and estimated marginal means with 95% confidence intervals were calculated using the emmeans package (Lenth *et al*., 2023).

A prerequisite for comparing ΔF and peak discharge rate between groups is that the number of identified MUs should not differ between the groups. Therefore, the effects of physical training experience and muscle on the number of test MUs identified were assessed using a generalised linear model with a log-link and Poisson error distribution, using a scaling factor (quasi-Poisson) to allow for overdispersion. When a significant effect was observed, Tukey post hoc correction was adopted for pairwise comparisons. The ratios of the number of test MUs identified between groups were calculated with 95% confidence intervals using the emmeans package.

Correlations between muscles regarding estimates of PIC magnitude were assessed by the Pearson correlation coefficient after averaging the data of the MUs for each muscle. The α level for all tests was 5%.

## RESULTS

### Participant characteristics

There were no significant differences between the groups in terms of age (F = 1.578, ω2 = 0.03, p = 0.221) and height (F = 1.451, ω2 = 0.02, p = 0.248). However, there was a significant difference in body mass (F = 5.208, ω2 = 0.19, p = 0.011), with the resistance group having a significantly higher body mass compared to the endurance group (+15.6 kg [3.7, 27.4], p = 0.008). No other between-group difference was found (all p ≥ 0.183). The duration of physical training per week also differed significantly between groups (F = 120.983, ω2 = 0.87, p < 0.0001). Specifically, the weekly duration of physical training was longer for the resistance than both the endurance (+196 min [88, 305], p = 0.0003) and the inactive group (+655 min [549, 761], p < 0.0001). Additionally, the endurance group had a longer physical training time compared to the inactive group (+459 min [352, 565], p < 0.0001). It is important to note that resistance training usually includes more rest periods compared to endurance training, which is important to consider when interpreting these differences.

There were significant differences between groups in terms of MVC for knee extension (F = 6.101, ω2 = 0.22, p = 0.006) and plantar flexion (F = 3.762, ω2 = 0.13, p = 0.034), but not for dorsiflexion (F = 1.700, ω2 = 0.04, p = 0.198). Post hoc pairwise comparisons revealed that the resistance group had significantly higher knee extension maximal torque compared to both the endurance (+77.0 Nm [18.0, 136.1], p = 0.008) and the inactive group (+67.6 Nm [8.6, 126.7], p = 0.022), with no difference between the endurance and the inactive group (+9.4 Nm [-49.6, 68.4], p = 0.920). The resistance group exhibited significantly greater plantar flexion maximal torque compared to the inactive group (39.5 [2.2, 76.8] Nm, p = 0.036) but not significantly higher than the endurance group (+31.3 [-6.0, 68.6], p = 0.114), with no significant difference between the endurance and the inactive group (+8.15 [-19.1, 45.4], p = 0.854).

### Identification of motor units

The number of identified and analysed MUs are provided in Table 2. A total of 2230 MUs were identified across all participants and muscles among which 1287 were paired and therefore used as test MUs to estimate PIC magnitude. We failed to identify enough MUs to estimate PIC magnitude for 13 conditions (i.e. 13 muscles from 9 participants), among which 7 came from the inactive group. There was no significant interaction between the group and muscle factors in terms of the number of test MUs identified (p = 0.941), and no significant group effect (p = 0.424). However, there was a significant muscle effect (p < 0.0001). Post hoc pairwise comparisons between muscles revealed that the number of test MUs was higher in TA compared to SOL (ratio = 1.6 [1.1, 2.2], p = 0.004), VL (ratio = 2.7 [1.8, 4.2], p < 0.0001), and VM (ratio = 3.2 [2.0, 5.0], p < 0.0001). The number of test MUs was also higher in GM compared to SOL (ratio = 1.8 [1.3, 2.5], p < 0.0001), VL (ratio = 3.1 [2.0, 4.7], p < 0.0001), and VM (ratio = 3.6 [2.3, 5.7], p < 0.0001). In addition, the number of test MUs was higher in SOL compared to VL (ratio = 1.7 [1.1, 2.7], p = 0.009) and VM (ratio = 2.0 [1.3, 3.3], p = 0.0005). There were no differences between TA and GM (ratio = 0.9 [0.7, 1.2], p = 0.79) and VL and VM (1.2 [0.7, 2.0], p = 0.922).

**Table 2:**
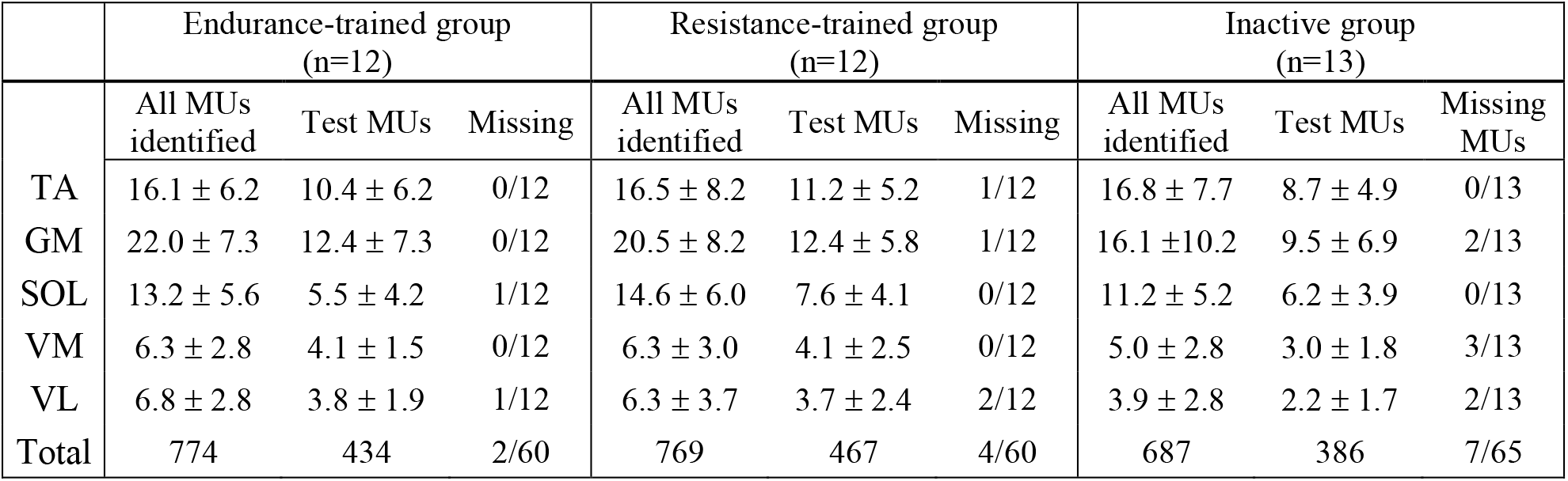
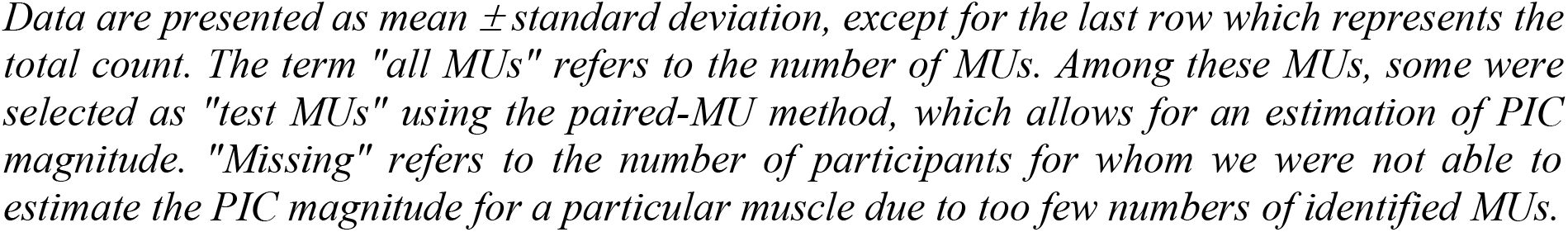
Number of identified and analyzed motor units.

|  | Endurance-trained group<br>(n=12) |  |  | Resistance-trained group<br>(n=12) |  |  | Inactive group<br>(n=13) |  |  |
| --- | --- | --- | --- | --- | --- | --- | --- | --- | --- |
|  | All MUs<br>identified | Test MUs | Missing | All MUs<br>identified | Test MUs | Missing | All MUs<br>identified | Test MUs | Missing<br>MUs |
| TA | 16.1 ± 6.2 | 10.4 ± 6.2 | 0/12 | 16.5 ± 8.2 | 11.2 ± 5.2 | 1/12 | 16.8 ± 7.7 | 8.7 ± 4.9 | 0/13 |
| GM | 22.0 ± 7.3 | 12.4 ± 7.3 | 0/12 | 20.5 ± 8.2 | 12.4 ± 5.8 | 1/12 | 16.1 ± 10.2 | 9.5 ± 6.9 | 2/13 |
| SOL | 13.2 ± 5.6 | 5.5 ± 4.2 | 1/12 | 14.6 ± 6.0 | 7.6 ± 4.1 | 0/12 | 11.2 ± 5.2 | 6.2 ± 3.9 | 0/13 |
| VM | 6.3 ± 2.8 | 4.1 ± 1.5 | 0/12 | 6.3 ± 3.0 | 4.1 ± 2.5 | 0/12 | 5.0 ± 2.8 | 3.0 ± 1.8 | 3/13 |
| VL | 6.8 ± 2.8 | 3.8 ± 1.9 | 1/12 | 6.3 ± 3.7 | 3.7 ± 2.4 | 2/12 | 3.9 ± 2.8 | 2.2 ± 1.7 | 2/13 |
| Total | 774 | 434 | 2/60 | 769 | 467 | 4/60 | 687 | 386 | 7/65 |

Data are presented as mean ± standard deviation, except for the last row which represents the total count. The term "all MUs" refers to the number of MUs. Among these MUs, some were selected as "test MUs" using the paired-MU method, which allows for an estimation of PIC magnitude. "Missing" refers to the number of participants for whom we were not able to estimate the PIC magnitude for a particular muscle due to too few numbers of identified MUs.

### Effect of physical training experience

#### Estimates of persistent inward currents (ΔF)

The mean ΔF values for each muscle and each group are depicted in Fig.2A. There was a significant main effect of muscle (TA: 4.7±1.2 pps, GM: 3.5±1.0 pps, SOL: 2.9±1.1 pps, VL: 1.8±1.0 pps, VM: 2.1±1.1 pps; F = 38.546, ω2 = 0.77, p < 0.0001), revealing differences between all muscles (all p ≤ 0.010), except between VL and VM (−0.22 [-0.62, 0.18] pps, p = 0.575) and between GM and SOL (0.55 [0.15, 0.94] pps, p = 0.055). There was no significant effect of physical training experience (RES: 3.1±1.4 pps, END: 3.2±1.6 pps, INA: 2.8±1.5 pps; F = 0.867, ω2 = - 0.0071, p = 0.429), nor a significant interaction between physical training experience and muscle (F = 1.037, ω2 = 0.0061, p = 0.426). This indicates that ΔF was not influenced by the physical training experience, irrespective of the muscle.

**Figure 1.**
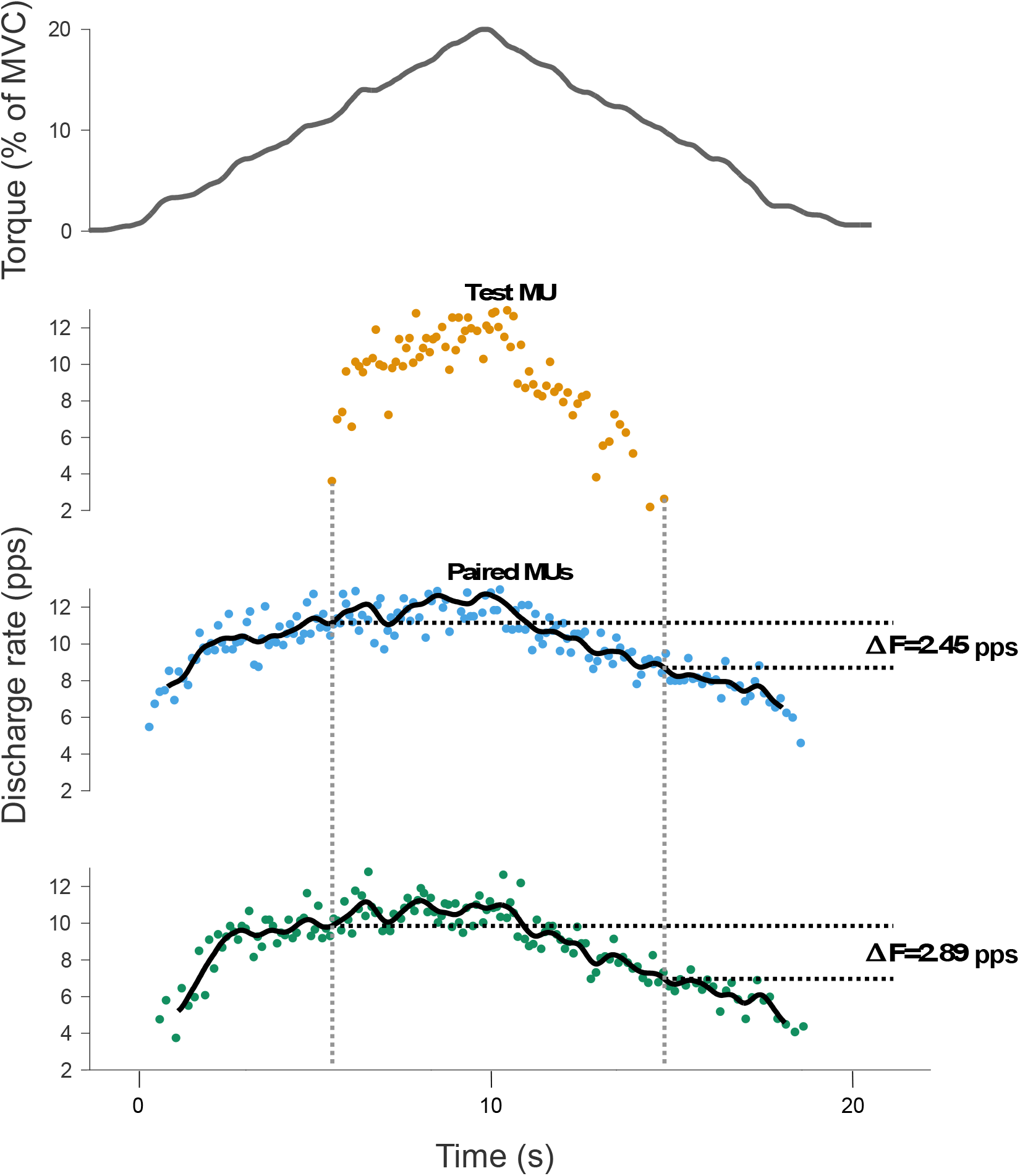
representative example of the ΔF calculation using paired motor unit analysis. The top panel shows the torque trace for a triangular contraction performed up to 20% of the participant’s maximal voluntary contraction torque. The subsequent panels display a test motor unit (orange color) and two control units (blue and green colors). The instantaneous discharge rate was smoothed with a 1-s Hanning window. The ΔF values obtained from the two control units were averaged, resulting in a single value for the test unit: (2.45+2.89)/2 = 2.67 pulses per second (pps).

**Figure 2.**
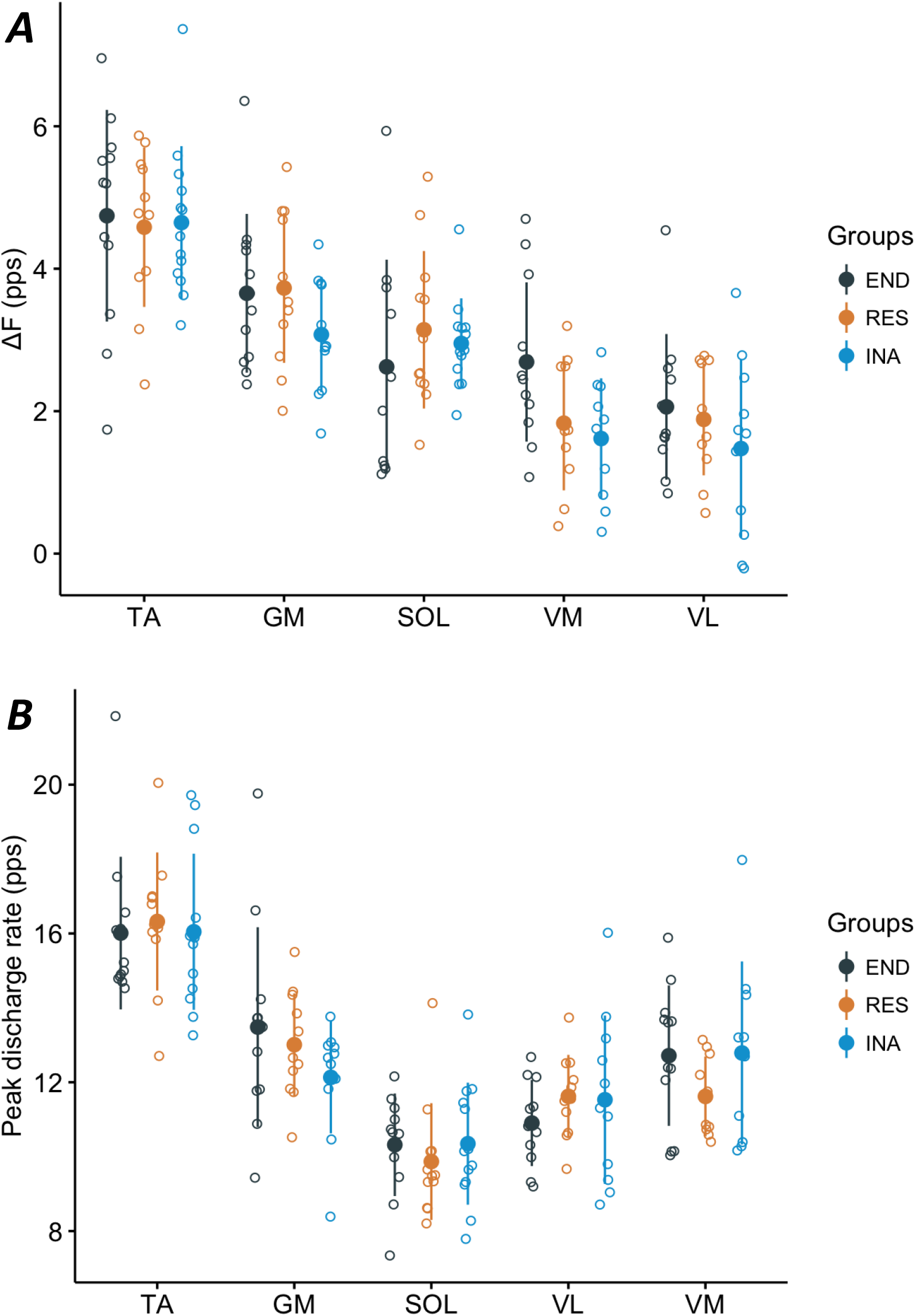
Estimates of persistent inward current magnitude (A) and peak discharge rate (B) according to physical training and muscles. The values were obtained from the tibialis anterior (TA), gastrocnemius medialis (GM), soleus (SOL), vastus medialis (VM) and vastus lateralis (VL). Circles are colored by groups: endurance-trained (END, black), resistance-trained (RES, orange) or inactive (INA, blue). Individual participant means are shown by open circles and group mean and SD by filled circles and bars, respectively. ΔF estimates the magnitude of persistent inward currents. pps = pulses per second. In A, there was a significant muscle effect, all muscles being significantly different except VL and VM and GM and SOL. However, no significant effect of physical training experience nor a significant interaction between physical training experience and muscle was found. In B, there was a significant muscle effect, all muscles being significantly different except VM and GM. However, no significant effect of physical training experience nor a significant interaction between physical training experience and muscle was found.

#### Peak discharge rate of motor units

The peak discharge rate values for each muscle are presented in Fig.2B. There was a significant main effect of muscle (TA: 16.1±2.0 pps, GM: 12.9±2.0 pps, SOL: 10.2±1.5 pps, VL: 11.4±1.6 pps, VM: 12.4±1.9 pps; F = 54.779, ω2 = 0.83, p < 0.0001), revealing differences between all muscles (all p ≤ 0.011), except between GM and VM (0.43 [-0.63, 1.48] pps, p = 0.766). However, there was no significant effect of physical training experience (RES: 12.5±2.6 pps, END: 12.8±2.8 pps, INA: 12.6±2.8 pps; F = 0.068, ω2 = -0.054, p = 0.935). There was a significant interaction between physical training experience and muscle (F = 2.216, ω2 = 0.17, p = 0.047). However, post hoc pairwise comparisons revealed no significant differences between groups, regardless of the muscle (all p ≥ 0.201). It indicates that peak discharge rate was not influenced by the physical training experience, irrespective of the muscle.

### Between-muscle correlations in estimates of persistent inward currents (ΔF)

We assessed the correlation of ΔF between muscles to assess specific associations between muscles (Table 3). There were significant correlations between VL and VM (r = 0.60 [95% CI 0.44, 0.85], p < 0.001), SOL and GM (r = 0.35 [CI 95% 0.012, 0.62], p = 0.041), and TA and GM (r = 0.49 [CI 95% 0.18, 0.71], p = 0.003). No significant correlation was observed for the other muscle pairs (all p ≥ 0.077).

**Table 3.** Pearson correlations of estimates of persistent inward current magnitude between muscles.

|  | VM | VL | TA | GM |
| --- | --- | --- | --- | --- |
| VL | **$r = 0.70$ [0.44 ; 0.85],**<br>**$p < 0.0001$** |  |  |  |
| TA | $r = 0.32$ [-0.04 ; 0.61],<br>$p = 0.077$ | $r = 0.19$ [-0.17 ; 0.51],<br>$p = 0.297$ | | |
| GM | $r = 0.20$ [-0.15 ; 0.52],<br>$p = 0.261$ | $r = 0.32$ [-0.04 ; 0.60],<br>$p = 0.078$ | **$r = 0.49$ [0.18 ; 0.71],**<br>**$p = 0.003$** | |
| SOL | $r = 0.12$ [-0.24 ; 0.45],<br>$p = 0.521$ | $r = 0.22$ [-0.12 ; 0.53],<br>$p = 0.208$ | $r = 0.14$ [-0.20 ; 0.45],<br>$p = 0.419$ | **$r = 0.35$ [0.02 ; 0.62],**<br>**$p = 0.041$** |

*Data presented are Pearson correlation coefficient between mean estimates of persistent inward current magnitudes, with 95% confidence interval and p-value. Statistically significant results are shown in bold.*

## DISCUSSION

In this study, we aimed to determine whether the estimates of α-motoneuronal PIC magnitude (ΔF) differ between young individuals with different physical training experience, i.e. resistance-trained, endurance-trained, and inactive. This study also aimed to determine whether the estimates of α-motoneuronal PIC magnitude are correlated across various lower limb muscles. Contrary to our hypothesis, the ΔF values did not differ across muscle groups irrespective of the muscle, and we only found significant correlations in ΔF value for three muscle pairs (out of 10). Overall, our findings suggest that i) the estimates of PIC magnitude in the lower limbs of young individuals are not influenced by physical training experience, and ii) neuromodulatory inputs are prone to muscle-specific and muscle group-specific regulations.

### No effect of training experience on estimates of PIC magnitude

Despite the substantial physical training experience differences among the groups, we found no differences in ΔF between groups, irrespective of the lower limb muscles. We also did not observe differences in peak discharge rate between groups for the five lower limb muscles assessed. It may be noted that the absence of differences in MU peak discharge rate does not necessarily imply an absence of differences in ΔF. For example, estimates of PICs can be larger in females compared to males despite a lack of sex-differences in peak discharge rates (Jenz *et al*., 2023). Since PICs significantly alter the intrinsic excitability of motoneurons (Heckman *et al*., 2009), an increase in PIC magnitude could represent a potential efficient adaptation that would allow for the generation of similar MU outputs with reduced descending ionotropic inputs. However, the results of this study suggest that despite the significant physical training practice of the participants in the endurance group (486 ± 114 min/week) and the resistance group (684 ± 150 min/week), no such adaptations occurred when compared to the inactive group (29 ± 29 min/week).

The absence of changes in α-motoneuronal PIC magnitude does not necessarily contradict the suggested increase in monoaminergic release with physical training in rat and mice (Ji *et al*., 2017; Ge & Dai, 2020). The effects of monoamines depend on the specific receptor subtypes present in the target neurons (Heckman *et al*., 2009). Thus, long-term physical training could potentially increase monoamine release without directly affecting PIC magnitude, while still contributing to an increase in intrinsic excitability. One possible mechanism that could contribute to enhanced motoneuron excitability is the downregulation of serotonin-mediated receptors, such as 5HT1A, involved in tonic inhibition. This downregulation has been suggested by changes in mRNA expression observed in rat models during endurance training (Woodrow *et al*., 2013; Perrier *et al*., 2018). Monoamines can also impact several other important α-motoneuron roperties that alter the intrinsic excitability of α- motoneurons (Powers & Binder, 2001), for example decreasing the spike afterhyperpolarisation amplitude (Berger *et al*., 1992; Manuel *et al*., 2006). These mechanisms may contribute to enhanced motoneuron excitability without change in PIC magnitude could also explain the differences between the results of this study and animal studies that report an increase in the intrinsic excitability of α-motoneurons with both resistance and endurance training (Krutki *et al*., 2017; MacDonell & Gardiner, 2018). Unfortunately, the paired-MU technique used in this study does not provide direct insight into monoaminergic inputs or changes in other intrinsic properties than PICs, as monoaminergic receptor sensitivity or ion channel function (Gorassini *et al*., 2002).

The absence of difference between the resistance and the inactive groups seems to be in contradiction with an increase in the estimates of PIC magnitude observed in older individuals after 6 weeks of resistance training (Orssatto *et al*., 2023). However, it is important to note that the estimates of PIC magnitude are typically reduced in the elderly (Hassan *et al*., 2021; Orssatto, Borg *et al*., 2021), which may be attributed to the age-related deterioration of the monoaminergic system (Shibata *et al*., 2006; Liu *et al*., 2020). Even though chronic physical activity may increase the secretion of monoamines (Ji *et al*., 2017; Ge & Dai, 2020), it is possible that even inactive young individuals had sufficient concentrations of monoamines to achieve comparable PIC magnitudes to trained individuals. In other words, the absence of difference of PIC magnitude between inactive young individuals and age-matched trained populations could be explained by a ceiling effect.

### Relationship in estimates of PIC magnitude between muscles

Contrary to our hypothesis, estimates of PIC magnitude were correlated for only 3 pairs of muscles (out of 10) (VL-VM, SOL-GM, and TA-GM). As magnitude of PICs depends on the level of monoaminergic drive onto the motoneurons (Lee & Heckman, 2000*b*), the presence of significant correlations between some muscles can be primarily attributed to the diffuse descending monoaminergic inputs to the α-motoneurons (Heckman, Hyngstrom *et al*., 2008; Wei *et al*., 2014). However, the overall low correlations and the lack of significant correlations between most of the tested muscles rather suggests that the magnitude of α-motoneuronal PICs is substantially prone to muscle-specific regulations. Interestingly, significant between-muscle correlations of estimates of PICs magnitude were found for muscles of the same group (VL-VM from quadriceps, SOL-GM from triceps surae), suggesting these regulations vary less among muscles from the same group (i.e. muscle group-specific regulations). In other words, regulations of monoamines inputs could be more common among muscles from the same group. Previous work in both animal (Kuo *et al*., 2003; Hyngstrom *et al*., 2007) and human (Mesquita *et al*., 2022) models have shown that inhibitory synaptic inputs can substantially reduce PICs activity. Acting as local regulators of PICs, the inhibitory inputs may explain muscle and muscle group-specific regulations. Notably, the inhibitory effect of Renshaw cells has been demonstrated to effectively deactivate PICs (Hultborn *et al*., 2003; Bui *et al*., 2008). Since the inhibitory effect of Renshaw cells extends to neighboring motoneurons (Ryall, 1970; Baldissera *et al*., 1981), it is plausible that it would result in concurrent deactivation of PICs in muscle from the same group. Irrespective of the specific mechanism, the explanation of muscle group-specific regulations by inhibitory inputs aligns with the proposed functional role of these inputs. Indeed, it is assumed that inhibitory inputs interact with the diffuse descending monoaminergic system to effectively control the intrinsic excitability of motoneurons (Heckman *et al*., 2009).

PICs are influenced by both noradrenaline and serotonin inputs, with noradrenaline being more influenced by arousal state and serotonin being increased along with motor output (Heckman *et al*., 2009). Thus, it should be emphasised that PIC magnitude has been estimated during three different tasks. These latter could have generated different monoamines release, which may explain preferential correlation towards muscles of the same group. However, this is unlikely to explain our results as the contraction intensity was normalised to maximal voluntary contraction, ensuring that each task elicited the same relative effort. In this way, it is important to note that a significant correlation was observed between TA and GM, which were assessed during different contractions. In addition, participants were deliberately kept in a calm state during all of the assessments, which likely led to similar noradrenaline release across tasks.

### Methodological considerations

Our study provides insights into the effects of different types of physical training experience on the estimates of PIC magnitude. We employed a cross-sectional design, allowing us to compare individuals with substantial practice in resistance or endurance training. Importantly, the comparison of PIC magnitude between groups has been based on a similar number of test MUs for all muscles. In addition, we assessed PIC magnitude in five different lower limb muscles, which is a larger number compared to previous studies in this area (Hassan *et al*., 2021; Orssatto, Mackay *et al*., 2021, 2023; Goodlich *et al*., 2023). This approach enabled us to provide consistent findings across multiple muscles and explore the relationship of PIC magnitude between muscles.

Estimates of PIC magnitude from this study are restricted to MUs recruited during low-intensity contractions, specifically between 0 and 20% of peak torque. This was due to the challenge of accurately measuring PIC magnitude at higher contraction intensities. Notably, the calculation of ΔF relies on low threshold MUs that provide a more accurate representation of synaptic inputs to the motoneuron pool given the more linear and proportional relationship of firing rate to synaptic input in the tertiary range (i.e. delayed firing) (Gorassini *et al*., 2002; Afsharipour *et al*., 2020). However, at higher contraction intensities, it becomes difficult to separate these low threshold MUs from larger, higher threshold MUs in the HDsEMG signal (Holobar *et al*., 2014). Additionally, the slow increase and decrease of force inherent to ramped contractions used in the paired MU analysis may induce fatigue when higher intensity contractions are performed, leading to confounding factors in PIC magnitude estimations. As a result, obtaining valid estimates of PIC magnitude values at higher contraction intensities remains uncertain. Therefore, the findings of this study cannot be directly extrapolated to higher-threshold MUs at higher contraction intensities. However, it is worth speculating that higher-threshold MUs at higher contraction intensities may exhibit similar activity (i.e. no differences in ΔF), despite differences in physical training experience. This speculation is notably supported by previous research showing that the administration of a competitive antagonist for the 5-HT2 receptor, which contributes to PICs, led to a reduction in MU firing rate irrespective of the contraction intensity. The reduction in firing rate was observed even at intensities of up to 70% of MVC, suggesting that the effects of serotonin on alpha-motoneurons are not dependent on the intensity of the voluntary muscle contraction being performed (Goodlich *et al*., 2023).

Regarding the statistical power of this study, the sample size was determined to be able to detect a difference of 0.8 pps in ΔF between groups. It is possible that the study was not adequately powered to detect smaller differences between the groups, although such differences might be of lower physiological relevance. However, it is worth noting that the effect sizes observed in this study were very low when examining the impact of physical training experience on estimates of PICs. Importantly, we performed a complementary analysis where both the resistance- and endurance-trained individuals were pooled in the same ‘trained’ group (n=24) and then compared to the inactive group. No significant differences were found (main effect group, p = 0.210; interaction group*muscle, p = 0.401), despite the larger sample size for this comparison. It makes us confident that the absence of between-group difference was not due to the relatively small sample size.

It should also be emphasised that results are limited to healthy young males in lower limb muscles, and thus cannot be generalized to other limbs nor populations, such as female and the elderly who may exhibit different PIC activity, as found in previous studies (Orssatto, Borg *et al*., 2021; Jenz *et al*., 2023).

## Conclusion

This study provides evidence that the estimates of α-motoneuronal PIC magnitude in MUs do not differ among young individuals who are resistance-trained, endurance-trained, or inactive during submaximal contractions of lower limb muscles. Additionally, the study suggests substantial muscle-specific regulations of the diffuse descending monoamine inputs, even though these regulations appear to be more common among muscles from the same group. Further investigations are needed to explore other potential factors that may contribute to the altered intrinsic excitability of motoneurons in response to various types of physical training, as suggested by animal studies. Additionally, further studies are required to investigate the mechanisms underlying the muscle-group specific regulations of motoneuronal PICs.

